# Co-occurring Amino Acid Substitutions Reveal Shared Evolutionary Links between Mammary Gland Location and Litter Size in Mammals

**DOI:** 10.64898/2026.01.25.701627

**Authors:** Aayan Behura, Ramyaa Manoharan, Susanta K. Behura

## Abstract

The mammary gland plays a critical role in mammalian development by producing milk to nourish offspring. The number and location of mammary glands vary among mammals. In humans and other primates that typically produce a single offspring, mammary glands are confined to the thoracic region. In contrast, litter bearing species possess mammary glands distributed along the milk line extending from the inguinal to the thoracic regions. In this study, we test the hypothesis that mammary gland location is evolutionarily linked to litter bearing capacity in mammals by performing large scale comparative and evolutionary analyses. We applied trait phylogeny Bayesian modeling to infer the coevolution of litter bearing capacity and mammary gland location (hereafter referred to as traits) and to assess the role of natural selection in shaping genes associated with this coevolution. To evaluate the functional relevance of the candidate genes, we conducted gene ontology and pathway enrichment analyses, inferred network and cluster patterns, and examined expression patterns in mammary glands and placentae. Additionally, we analyzed within species variation in protein sequences among pig breeds with low or high teat numbers and litter sizes to model the one half rule. Our results indicate that mammary gland location and litter bearing capacity are evolutionarily linked via site specific substitutions of amino acids, natural selection, and interconnected networks of a suite of proteins that are regulated in both mammary gland and placenta, and associated with specific biological functions including signal transduction, cell communication, and immune system function.

## Introduction

Animals adapt numerous strategies to reproduce and develop (1). Matrotrophy, or the provisioning of maternal resources to the offspring during fetal and postnatal development, is one of them (2–4). Recent studies suggest that phylogenetically unrelated species express a similar set of genes in the placenta and mammary gland to reproduce and develop (5). Though mammary glands are indispensable for the survival of all mammals (6), they show striking variation in the location and number among species (7). In humans and primates, they are located in the chest (thoracic) regions while in ruminants and species that produce litters, they are located in the inguinal, abdominal, thoracic and often cervix regions (8). In rare cases, they are also located in the axillae and thighs (9, 10).

The position of the mammary gland is thought to confer evolutionary benefits for species (11). For instance, in humans, the positioning of the mammary gland has allowed for breastfeeding to align with humans’ bipedal posture, an advantage which has been key to infant survival throughout our evolution (12). The molecular mechanisms of how mammary buds are developed at different locations are not clear, but it is known that the epithelial-mesenchymal interactions play crucial roles in mammary bud development to guide the formation, growth, and branching of the mammary glands (13). Besides location, the number of teats also play a key role in mammalian reproduction and development (14–16). The "one-half rule" about the teat number states that mean litter size is typically one-half the number of mammary glands (17). Broadly, species with more nipples clearly can support larger litters; for example, rodents have 10–12 nipples and tend to have litters of 6–14 pups, while those with two nipples or udders, like primates and ruminants, typically bear fewer offspring (18). In pigs, it was found that 14 or more teats increased litter size at birth compared to just 11-13 teats (19). In rabbits, a close relationship exists between teat number and large litter size (14). To relate the number of teats and their location, past work has developed a "teat formula" which expresses the total number of axillary, thoracic, abdominal, and inguinal teats for any given species (15).

Overall, single offspring-producing species tend to have a single thoracic pair while those with large litters tend to have multiple pairs extending along the entire length of the milk lines (20, 21). However, the molecular basis underlying these evolutionary patterns remains poorly understood. In this study, we hypothesize that mammary gland location and litter-bearing capacity are evolutionarily linked. To test this hypothesis, we performed large-scale comparative and evolutionary analyses of protein sequences among 18 mammalian species. We identified fixed changes in proteins by comparing the multiple sequence alignments of proteins among species with thoracic or non-thoracic location of mammary glands, and species with litter-bearing or single-offspring capacity. Next, we performed Bayesian modeling to infer co-evolution between litter production capacity and mammary gland location, and investigated functional relevance of the associated genes using gene ontology and pathway enrichment analyses, as well as network and cluster analyses. Finally, we examined patterns across pig breeds that differ in litter size and teat number, and assessed the expression of candidate genes in the mammary gland and embryonic mammary buds in mice. The results of this study show that the evolutionary links between mammary gland location and litter bearing capacity in mammals are associated with fixed changes in amino acids of specific proteins that play key roles in mammary gland development, signal transduction, cell-cell communication, and immune system function.

## Materials and Methods

### Species selection, and extraction of sequence data

In this study, a total of 18 mammalian species were used that varied in mammary gland location, number and litter-bearing capacity (**Table 1**). These species are phylogenetically diverse with divergence times that vary from 7 million years ago for the human-chimpanzee split (22) to roughly 99 million years ago for elephant and uranotherian mammals (23). The *TimeTree* (24) annotation and the plot of evolutionary relationships and divergence times among the 18 species are shown in **Figure 1A**. The species are grouped based on the mammary gland location and litter-bearing capacity, here referred to as ‘traits’. Henceforth, the species groups are abbreviated as either L or NL to represent litter or non-litter form of reproduction, and as T or NT to represent the thoracic or non-thoracic location of mammary glands. To investigate the molecular signatures associated with the L/NL or T/NT traits, we compared the amino acid sequences downloaded for each of the 18 species from Ensembl. The assembly versions of the species are listed in the supplementary data **Text S1**. We wrote custom scripts (**Text S2**) that used the downloaded amino acid sequence files in FASTA formats to extract sequences of the proteins commonly present across the 18 species (n=6,002).

**Figure 1.**
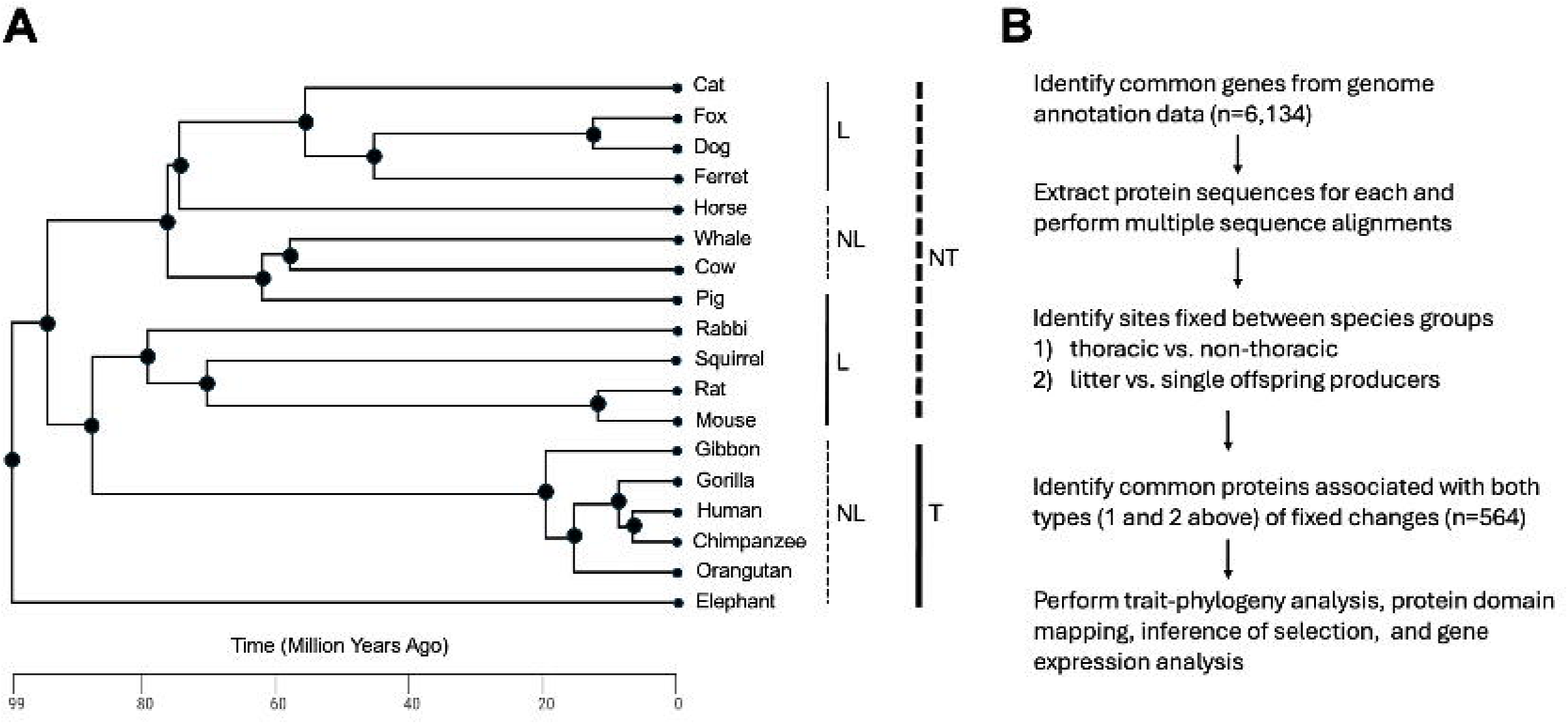
Species selection and comparative framework. (A) Phylogenetic relationships of the 18 mammalian species included in this study, categorized by litter-producing capacity (litter vs. non-litter) and mammary gland location (thoracic vs. non-thoracic). (B) Schematic overview of the computational pipeline used to extract orthologous protein sequences, perform multiple sequence alignments, identify fixed amino acid substitutions distinguishing L/NL and T/NT species, and conduct downstream evolutionary and functional analyses.

**Table 1.** Animal species used in the study. The reproductive and mammary related metadata are included.

| **Animal** | **Reproduction** | **Mammary**<br>**location** | **Teat**<br>**count** |
| --- | --- | --- | --- |
| Whale | Non-litter | Non-thorasic | 2 |
| Horse | Non-litter | Non-thorasic | 2 |
| Gorilla | Non-litter | Thorasic | 2 |
| Human | Non-litter | Thorasic | 2 |
| Elephant | Non-litter | Thorasic | 2 |
| Gibbon | Non-litter | Thorasic | 2 |
| Chimpanzee | Non-litter | Thorasic | 2 |
| Orangutan | Non-litter | Thorasic | 2 |
| Cow | Non-litter | Non-thorasic | 4 |
| Cat | Litter | Non-thorasic | 8 |
| Ferret | Litter | Non-thorasic | 8 |
| Squirrel | Litter | Non-thorasic | 8 |
| Dog | Litter | Non-thorasic | 10 |
| Mouse | Litter | Non-thorasic | 10 |
| Rabbit | Litter | Non-thorasic | 10 |
| Fox | Litter | Non-thorasic | 10 |
| Rat | Litter | Non-thorasic | 12 |
| Pig | Litter | Non-thorasic | 14 |

### Multiple sequence alignments and amino acid substitution analysis

We performed multiple sequence alignment for amino acid sequences of each of the common proteins among the 18 species using Clustal Omega (ClustalO) (25). We wrote a script (**Text S3**) to analyze individual sites in the alignments to identify amino acids *fixed* or *nearly fixed* between L vs. NL or T vs. NT groups. We define a *fixed site* as a position in a multiple sequence alignment where the same amino acid is present in all species of one group and a different amino acid is present in all species of the other group. A *nearly fixed site* is defined as a position where the same amino acid is present in all but one species of one group and a different amino acid is present in all or all but one species of the other group. The amino acids representing the fixed/ nearly fixed variant sites between L and NL species were referred to as L/NL substitution; and those between T and NT species were referred to as T/NT substitutions. The number of L/NL and T/NT substitutions was determined for each protein separately for the L, NL, T and NT groups. Next, we performed hierarchical clustering of the number of substitution sites among the 20 amino acids using Ward’s method (26). The dendrograms representing the cluster patterns of amino acids between L vs. NL and T vs. NT were generated using the R package Dendextend (27).

### Trait-phylogeny modeling and evolutionary analysis

The gene sequences coding for the proteins associated with the L/NL and T/NT substitutions were subjected to phylogenetic analysis. Using the PAL2NAL tool (28), multiple sequence alignment was performed on the coding DNA sequences of each gene from the 18 species (downloaded from the Ensembl Genome Browser). PAL2NAL performs codon alignment of genes guided by the alignment of the amino acid sequences of the corresponding proteins. The aligned output sequences were saved in PAML format (29) as well as FASTA format (30). The PAML formatted sequence files were used for calculation of ratio of non-synonymous vs. synonymous changes (dN/dS) in each gene in a pair-wise manner between species using the CODEML tool within PAML (29). PAML is a collection of tools to perform evolutionary analysis of DNA or protein sequences based on maximum likelihood methods (31). To run CODEML in batch, control files listing the sequence file and run parameters were programmatically generated for each gene. The run parameters included: model = 0, NSsites = 0, runmode = −2, and clock = 0. CODEML was run in batch using a for-loop command to process each gene and generate the pair-wise dN/dS ratios. These ratios were parsed into two columns (species-pair compared and dN/dS ratio) using AWK.

The FASTA aligned sequences of the genes were used to generate neighbor-joining phylogenetic trees in nexus format with the tree root placed at the same species (whale) for each tree using the Linux-based phylogenetic tool Phyx (32). A for-loop bash command was used to create the phylogenies of all the genes using Phyx. The BayesTraits V4 tool (33) was then used to analyze the phylogenetic trees along with the species trait data (i.e. location of mammary gland and litter-producing capacity, see **Table 1**) for performing trait-phylogeny modeling. The trait data of the species were coded in binary forms. Litter production was coded as 1 (produces litters) or 0 (single offspring), and mammary gland location was coded as 0 (thoracic only) or 1 (non-thoracic or multiple locations). The trait correlation analysis was performed by comparing the likelihood of a discrete independent model with the likelihood of a discrete dependent model. The independent model assumes that the mammary gland’s location evolved independent of the evolution of litter production capacity while the discrete dependent model assumes that they evolved in a correlated manner. The model likelihoods were estimated using Markov chain Monte Carlo method (34) with 100 stones and 1000 iterations per stone. The priors were set to an exponential with a mean of 10 in each. The likelihood ratio of two models was compared by calculating the Log Bayes Factor (LBF) by subtracting the marginal likelihood estimate of the independent model from that of the dependent model and multiplying by two (35).

### Mapping variant sites to conserved protein domains

To test whether protein regions accumulating L/NL and T/NT substitutions were associated with specific functional domains, we analyzed flanking sequences around variant sites. For each protein, we extracted 25 amino acids on both sides of each variant site using the *substring* bash command. An amino acid sequence length of 50 was selected because a majority of the conserved domains in protein sequences have a minimum threshold length of 50 (36). The extracted sequences were mapped to the protein conserved domain database (37) via PSI-BLAST (38) to identify if they are located in specific domains.

### Network analysis and key player prediction

Network analysis of the variant amino acids was performed using the mutual information (MI) method (39). MI is a concept in information theory to quantify the amount of information shared between two random variables. It measures how much knowing one variable reduces uncertainty about the other. We determined the pair-wise MI scores between amino acids based on the variation of amino acid count data. Network analysis was performed using the R package minet (39). Centrality test (40) was performed to predict the key players that played central roles within each network.

### Gene ontology and pathway enrichment analysis

Gene ontology (GO) enrichment analysis was performed to infer the function of the proteins associated with the L/NL and T/NT substitutions using the *Panther* GO enrichment tool (41). The significance (False Discovery Rate) of enrichment was determined by the Fisher exact test. Pathway enrichment analysis was performed by using the *Reactome* database (42). The significance of pathway enrichment by hypergeometric test and data visualization were performed using the tools implemented within the *Reactome*.

### Analysis of gene expression in cell types using Single-Cell Portal

The expression of selected genes coding for the proteins associated with the observed amino acid substitutions was assessed in different cell types of developing mammary gland in mice using the Single-Cell Portal (SCP) database and tools (43). SCP allows researchers to analyze gene expression across cell types through interactive tools, including differential expression analysis, finding marker genes of different cell clusters, and visualization of gene expression patterns in single cells. We used the single-cell RNA expression data of mouse embryonic mammary gland (44) included in SCP to query selected genes identified from the current study and determined the expression differences in different cell types of developing mammary gland at a single-cell resolution.

### Analysis of mammae number and litter size in pig breeds

In this analysis, we selected specific pig breeds such as Tibetan, Rongchang, Bamei, Large White, Berkshire and Hampshire that have relatively fewer number of teats compared to the number of teats in Pietrain, Jinhua, Landrace, Wuzhishan, Usmarc and Meishan breeds. The amino acid sequences of the annotated proteins from these breeds were downloaded from Ensembl. The sequences of each protein were aligned among the 12 breeds and variant amino acids fixed or nearly fixed between the two breed groups were identified using the approach described above.

### Analysis of placental expression of genes that are associated with N/NL and T/NT substitutions in mammary gland

We analyzed gene expression of term placenta of three species (*Homo sapiens, Pan paniscus and Loxodonta Africana*) that have only two mammae compared to three species (*Mus musculus, Monodelphis domestica, Canis familiaris*) that have multiple mammae. The gene expression data of term placenta of these species were generated from a previous study (45). We extracted the placental expression data of the six species for genes coding for the proteins associated with the L/LN and T/NT substitutions identified in the present study. The data were merged by genes across the six species, and subjected to differential expression analysis between the two species groups using the R package *edgeR* (46).

## Results

### Amino acid substitutions in specific proteins distinguish L/NL and T/NT species

In this study, we analyzed the variation of amino acid sequences in individual proteins (n=6,002) among 18 phylogenetically diverse mammals (**Figure 1A**). The species were selected based on their differences in the location of mammary glands and litter-producing capacity (**Table 1**). The 6,002 proteins are commonly present across the selected species based on orthology annotation by Ensembl (47). The data analysis approach is illustrated in **Figure 1B**. We developed a computational pipeline to extract amino acid sequences of the 6,002 proteins across the 18 species, performed multiple sequence alignment and compared the individual sites in each alignment between the litter (L) producing vs non-litter (NL) producing species as well as between species with thoracic (T) vs non-thoracic (NT) locations. The scripts are provided in the supplementary data (**Text S1**). The analysis identified specific proteins (n=533) associated with either fixed or nearly fixed substitutions of amino acids (see Methods for definition) between L and NL species as well as between T and NT species (**Table S1**). Both types of substitutions, referred to as L/NL and T/NT, co-occurred in each of these proteins. The distance between the L/NL and T/NT substitutions varied among the proteins. To assess the distribution patterns of these substitutions relative to their distances, we made equal-length intervals (10 amino acids) of the distances and calculated the number of substitutions associated with each interval. The analysis showed that the number of substitutions decreases as the distances increase to 100 or more amino acids (**Figure S2**). All the 20 amino acids were represented by the substitutions (**Table 2**). However, they showed different cluster patterns in different species groups based on the number of the variations they represented in the individual proteins (**Figure 2**). The plot in **Figure 2** shows a side-by-side comparison of dendrograms representing the hierarchical cluster patterns of the individual amino acids between L vs. NL and T vs. NT species. It shows that leucine and alanine cluster into a separate group in the L species but group with several other amino acids in the NL species. Similarly, it was observed that alanine and threonine cluster into a separate group in the T species but group with other amino acids in the NT species. These contrasting cluster patterns suggested that the fixed changes that co-occurred in these proteins are associated with the amino acids in a non-random manner thereby biasing the substitutions with specific amino acid(s) among the species groups. To further understand the relationships of the biased patterns of amino acid substitutions among the four species groups (L, NL, T and NT), we determined the mutual information (MI) scores among the four groups in a pairwise manner. **Figure 3A** shows a mosaic plot representing the relationships among the species groups based on the MIs calculated from the variance of the number of fixed changes associated with each of the 20 amino acids across the 533 common proteins. **Figure 3B** shows the interconnectedness in the MI of amino acid variation in the form of a circos plot. It represents the visualization of how a species group is interconnected with the other three groups supported by the color-coded ribbons linking among the groups shown on the circumference. The cophenetic distance between cluster dendrograms and the cluster correlation patterns between species groups are shown in **Figure S3**.

**Figure 2.**
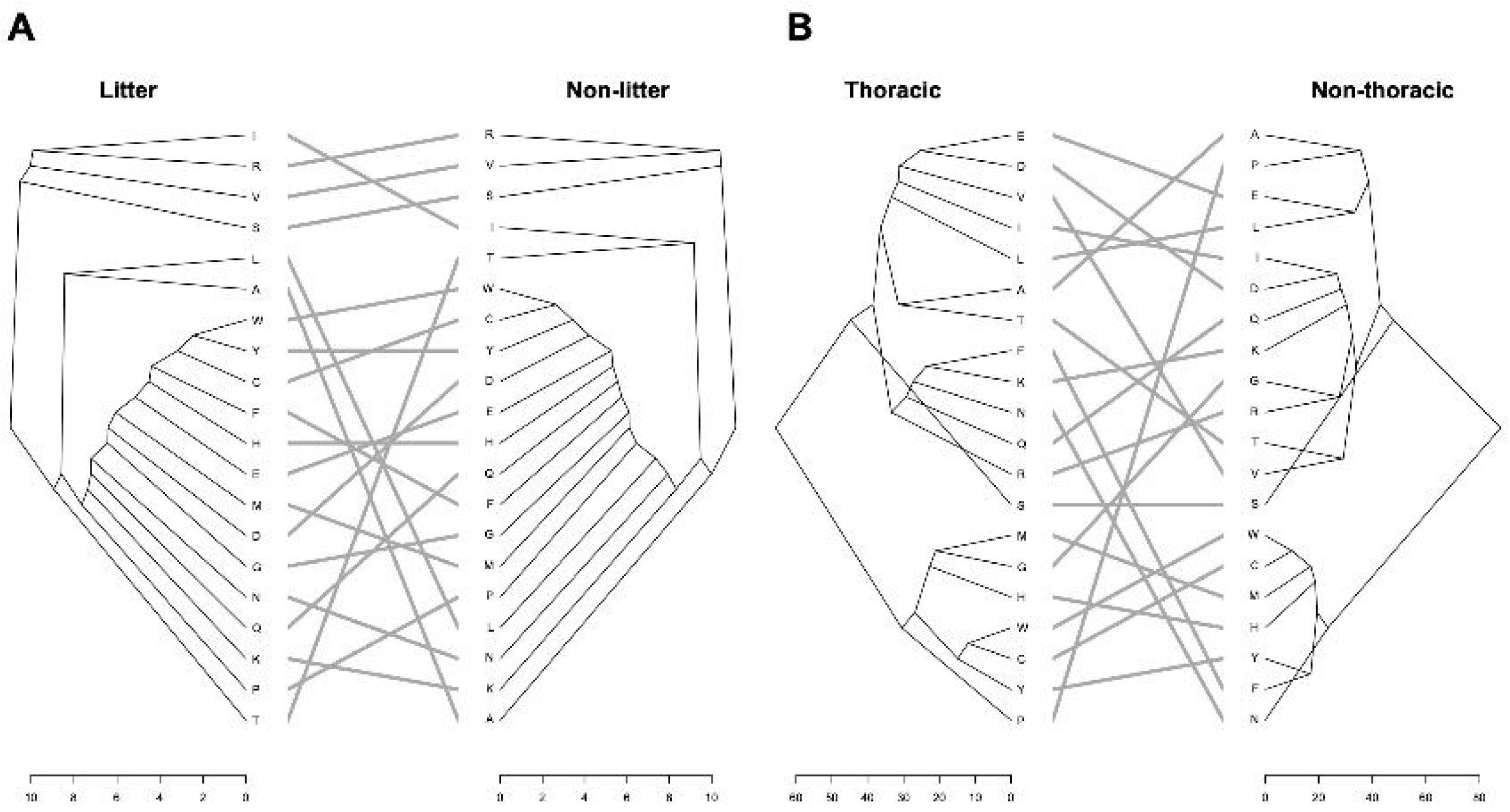
Amino acid substitution clustering patterns differ among species groups. Hierarchical clustering of the 20 amino acids based on the number of fixed substitutions observed in proteins distinguishing (left) litter-producing vs. non-litter-producing species and (right) thoracic vs. non-thoracic mammary gland location. Distinct clustering patterns indicate non-random biases in amino acid substitutions across species groups.

**Figure 3.**
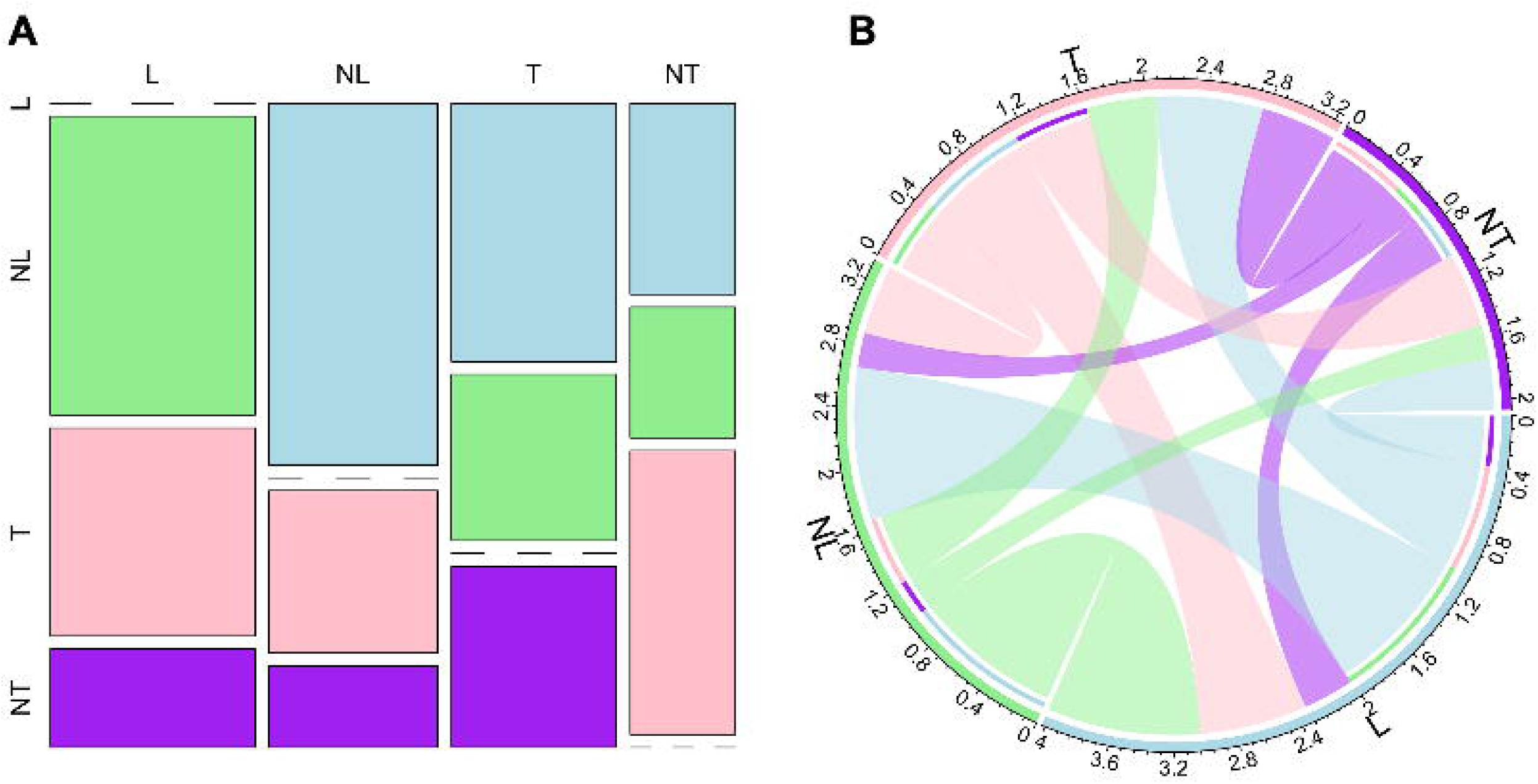
Mutual information reveals coordinated amino acid variation. (A) Mosaic plot showing pairwise mutual information (MI) scores among the four species groups (L, NL, T, NT), calculated from the variance in amino acid substitutions across the 533 shared proteins. (B) Circos plot visualizing interconnectedness among species groups, where ribbon thickness reflects the strength of MI, highlighting coordinated patterns of amino acid variation.

**Table 2.**
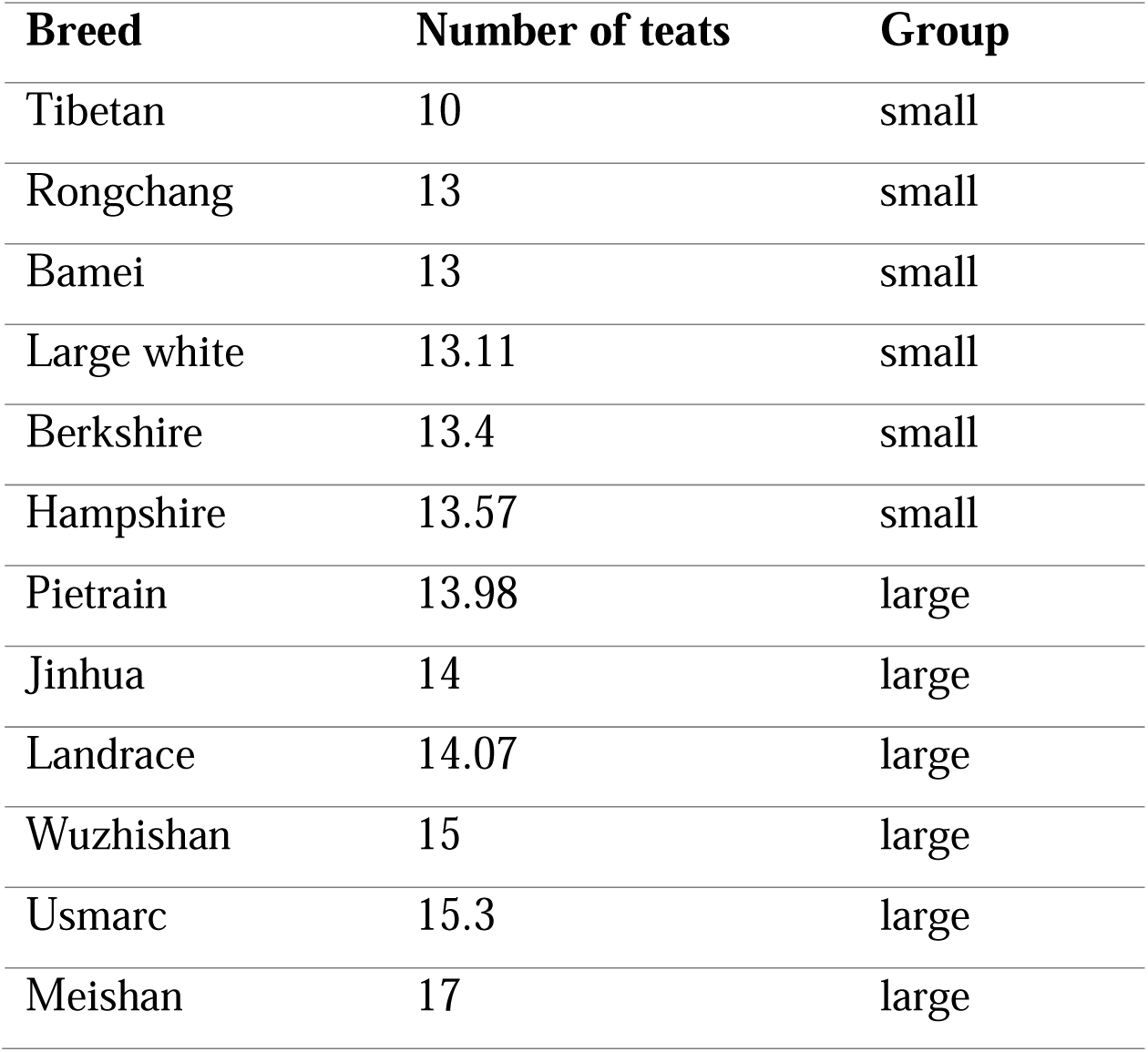
List of pig breeds used in the analysis. Teat counts in the breeds are shown.

| **Breed** | **Number of teats** | **Group** |
| --- | --- | --- |
| Tibetan | 10 | small |
| Rongchang | 13 | small |
| Bamei | 13 | small |
| Large white | 13.11 | small |
| Berkshire | 13.4 | small |
| Hampshire | 13.57 | small |
| Pietrain | 13.98 | large |
| Jinhua | 14 | large |
| Landrace | 14.07 | large |
| Wuzhishan | 15 | large |
| Usmarc | 15.3 | large |
| Meishan | 17 | large |

### Evidence for positive selection acting on genes associated with the variant proteins

The 533 genes coding for the proteins associated with the L/NL and T/NT variations, described above, were further analyzed to investigate evolutionary patterns using the CODEML tool in the Phylogenetic Analysis by Maximum Likelihood (PAML) software (29). We used CODEML to calculate the dN/dS ratio, where dN represents the rate of nonsynonymous mutations and dS represents the rate of synonymous mutations, for each gene. The results of this analysis showed that more than 95% of the genes coding for the proteins associated with the L/NL and T/NT variations (508 of the 533) were associated with either purifying or positive selection (Table S2). While a few genes (n=48) showed evidence for positive selection with dN/dS > 1, the majority of these genes (n=460) showed evidence for purifying selection with dN/dS < 0.2.

### Modeling evolutionary links between mammary gland location and litter producing capacity

To further infer evolutionary links between mammary gland location and litter producing capacity in mammalian species, we performed trait-phylogeny modeling (33). The trait data of the species was represented in binary forms (0 and 1) based on whether or not a species has mammary glands in the thoracic region only and also whether or not the species produces litter (**Table 1**). The trait data was analyzed with the phylogenetic trees of the genes (n=533) described above. The evolutionary tests were performed using the discrete trait modeling methods implemented in *BayesTraits* (33). In this analysis, rooted neighbor-joining phylogenetic trees (**Figure 4**) were analyzed to compare the likelihoods of the discrete independent and dependent models described in the Methods section. The likelihood estimates of both models are listed in **Table S3**. The comparison of likelihood estimates yielded a log Bayes Factor above 2 for the majority (529 out of 533) of the genes, supporting an evolutionary link between mammary gland location and litter production capability in mammalian species.

**Figure 4.**
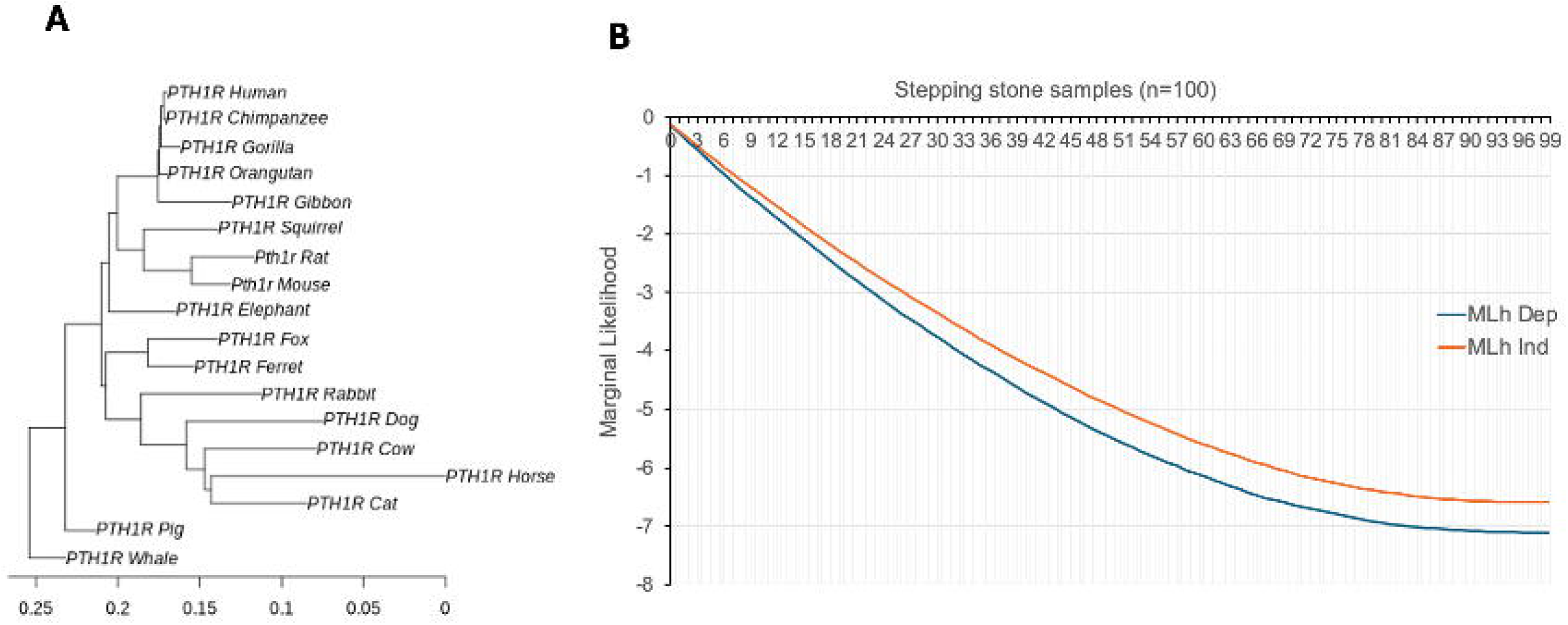
BayesTraits analysis supports correlated evolution of traits. (A) Representative rooted gene phylogeny used for discrete trait modeling, with the root placed at whale. (B) Distribution of log Bayes Factors comparing dependent and independent evolutionary models for mammary gland location and litter-producing capacity across 533 genes. Positive values indicate stronger support for correlated evolution.

### Variant amino acids in conserved domains, and patterns of protein network

We observed that specific amino acid substitutions were localized within conserved domains (CDs) of the proteins (**Table S4**). CDs are crucial functional and structural units, ranging from 50 to 250 amino acids in length, that dictate essential functions of proteins such as DNA binding, enzyme activity, and protein interactions across diverse species (37). We extracted amino acid sequences centering the observed substitution sites, with 25 residues on either side of the substitutions so that the subsequence length corresponds to the minimum size of CDs (50 amino acids), and then mapped those sequences to NCBI CD database (37) to identify the hits (**Table S4**). The analysis identified proteins (n=222) wherein the L/NL and T/NT co-occurred either in the same domain (n=152) or different domains (n=70) (**Table S5**). Thus, a majority of these proteins (68.5%) were associated with both types of substitutions in the same domains, suggesting that amino acid substitutions in the conserved domains may be differentially coordinated in different species groups. To further test this assumption, we applied a network analysis approach to identify the key proteins and their pattern of networks based on the number of amino acids that were associated with CDs. By performing MI network and key player analyses (39, 40), we observed that specific proteins played key roles in the coordinated changes of CDs among different species groups (**Figure 5**). Beyond that, we also observed that amino acid contexts in CDs (amino acids that are next to each other within a domain) also influences protein networks among the species (**Figure 6**).

**Figure 5.**
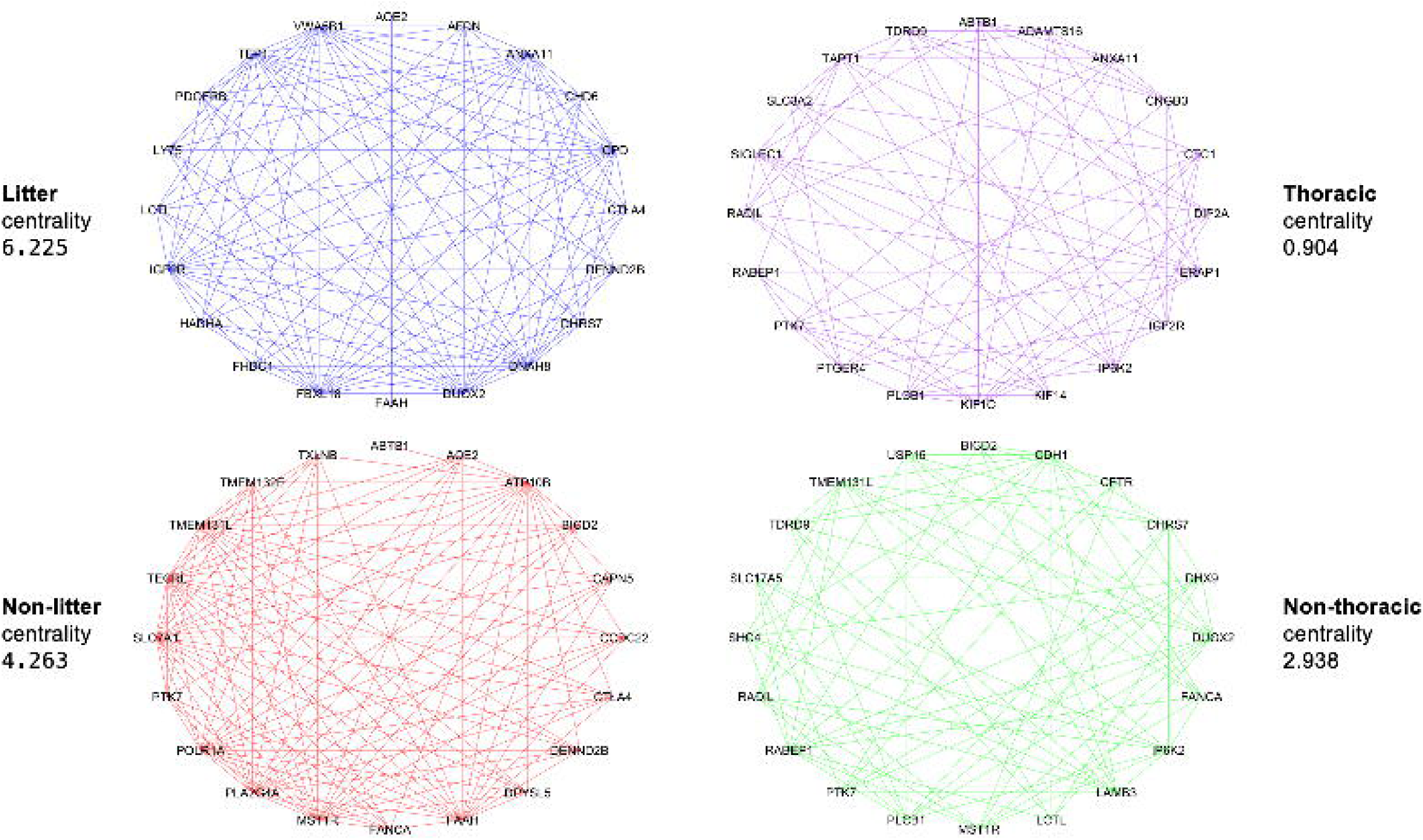
Network analysis of key proteins associated with variant amino acids. Mutual information–based protein interaction network constructed from conserved domain–associated amino acid substitutions. Nodes represents proteins, and edges indicate coordinated variation of the amino acids among proteins.

**Figure 6.**
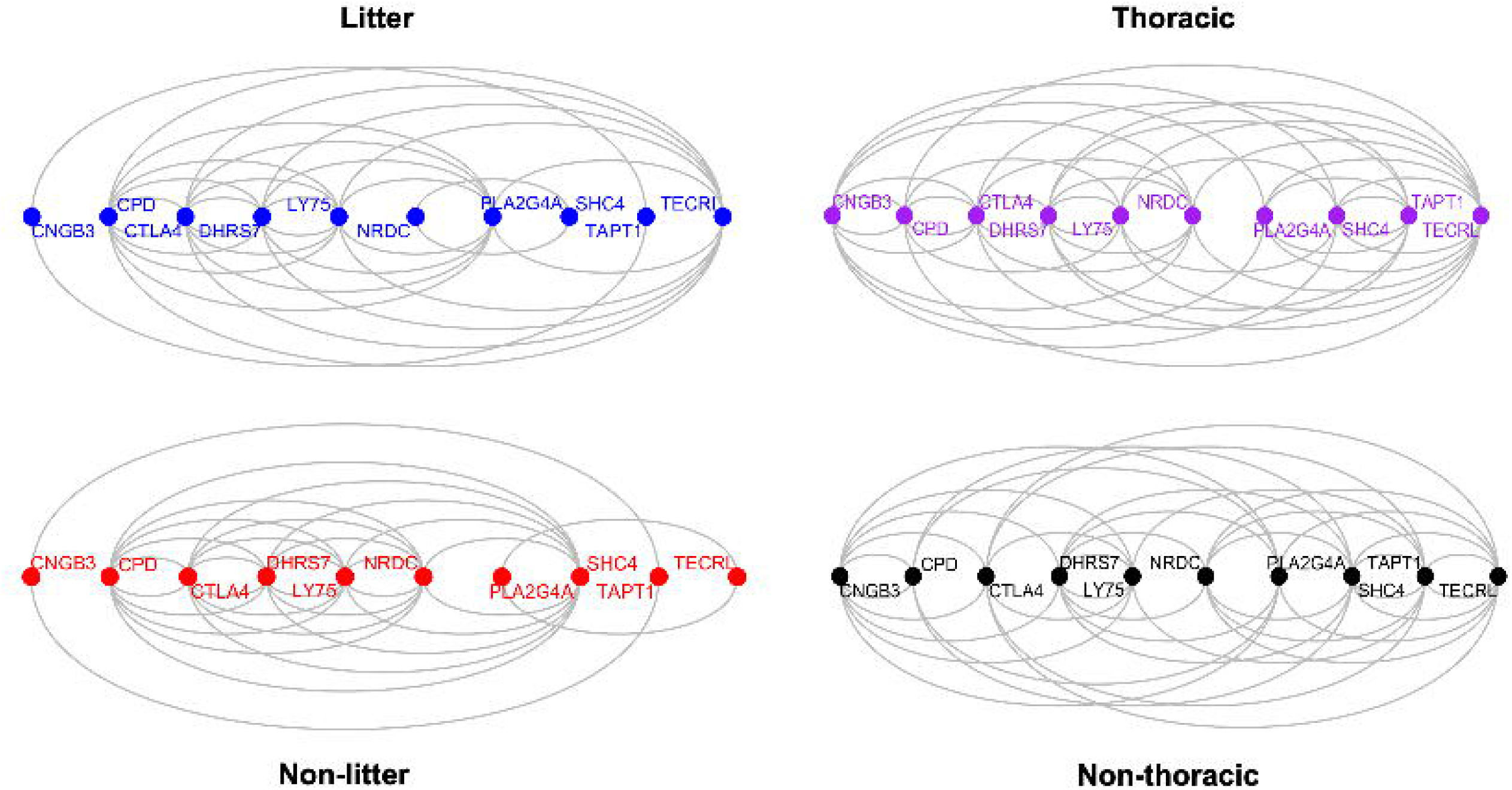
Amino acid context within conserved domains shapes protein networks. Network visualization of protein with variations of amino acid contexts within conserved domains.

### Functional annotation of the proteins associated with the amino acid substitutions

We performed functional annotation of the proteins (n=533) that were associated with the L/NL and T/NT amino acid substitutions. First, we performed gene ontology (GO) analysis to determine if specific biological functions are enriched among genes coding for these proteins (**Table S6**). The GO enrichment analysis showed a list of 39 biological functions that are significantly over-represented (False Discover Rate or FDR < 0.0005 and fold-enrichment > 1.5). The functions relating to sequestering of TGF beta in extracellular matrix, regulation of gap junction assembly, positive regulation of membrane protein ectodomain proteolysis, and amine transport emerged as the top GO terms that were enriched by more than 10-fold. We also performed pathway enrichment analysis with the same list of 533 proteins to determine if they were associated with specific pathways (**Figure 7**). The vesicle-mediated transport, membrane trafficking, interferon signaling, cell junction organization, and developmental cell lineages emerged as the top five enriched pathways from this analysis (**Table S7**). Moreover, pathways related to developmental lineages of the mammary gland cells including mammary gland luminal epithelial cells, myoepithelial cells, and stem cells were also significantly enriched.

**Figure 7.**
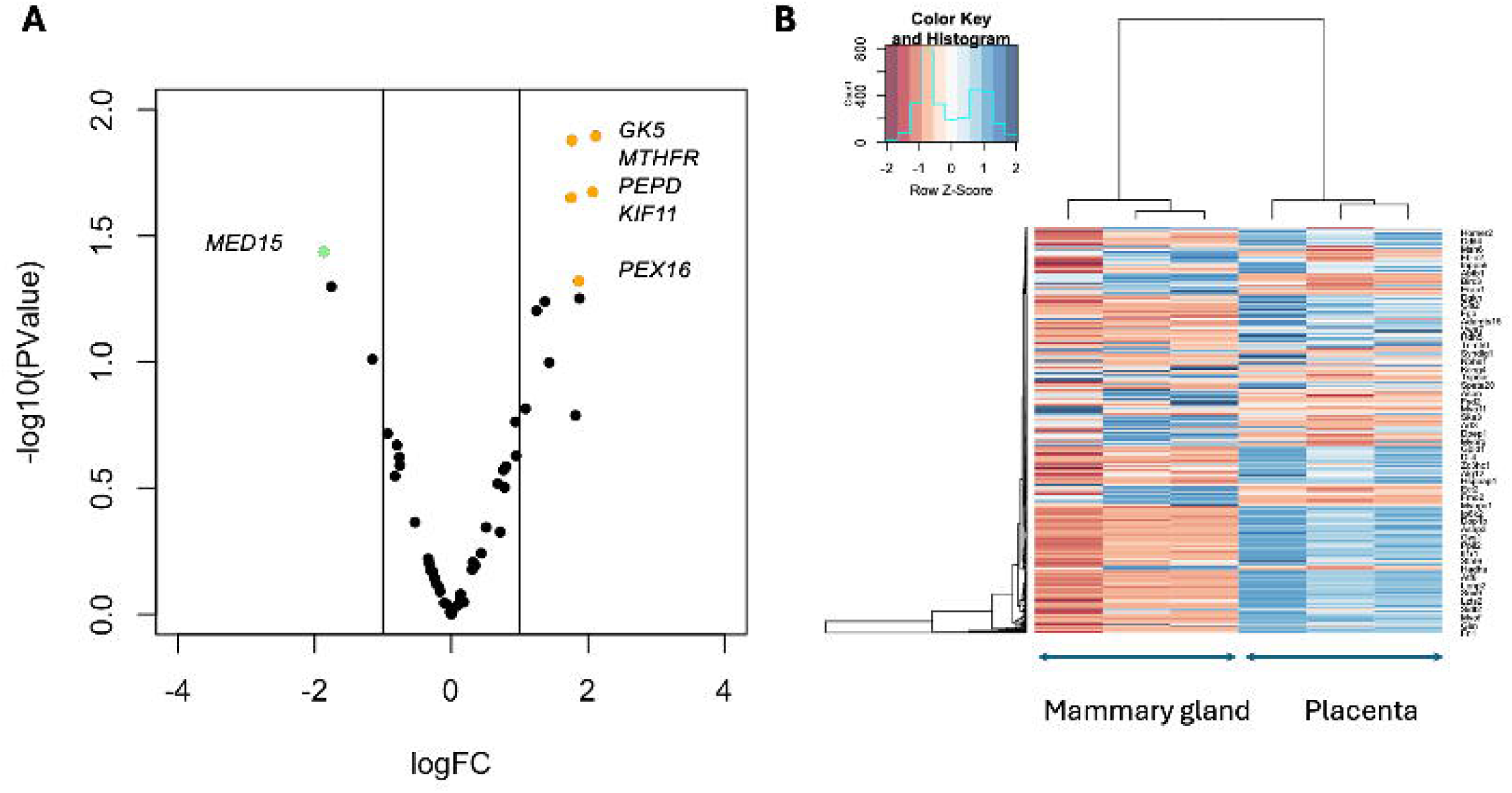
Pathway enrichment of proteins with L/NL and T/NT substitutions. Summary of significantly enriched pathways identified from pathway enrichment analysis of the 533 proteins. Pathways related to vesicle-mediated transport, membrane trafficking, cell junction organization, developmental signaling, and mammary gland cell lineages are prominently represented.

Previously, *Giraddi et al.* measured single cell RNA sequences and found 22,185 genes expressed during mammary bud development in mice (44). We investigated the expression level of the 533 genes using the Single-Cell Portal (43). The data is provided in **Table S8**. The plot in **Figure 8B** shows a representative plot for the expression pattern of the gene encoding for TAPT1 (transmembrane anterior posterior transformation protein 1) in different cell types of the embryonic mammary bud in mice.

**Figure 8.** Expression of TAPT1 in different cell types in the developing mammary gland of mice. Single-cell RNA-seq expression pattern of TAPT1 across cell types in the embryonic mammary bud, illustrating cell-type-specific expression during development. The UMAP plot shows the cell clusters are color coded and the number of cells in the cluster are shown. The scatter plots shown the relative expression level of each cell (each dot represents a cell). The scale on the right shows the color codes for expression levels.

### Relationship between mammary gland variation and litter size in pigs

To further analyze teat number variation within species, we performed compared analysis of protein sequences among twelve breeds of pigs (**Table 2**). Six of these breeds have an average teat count of 12.8, here classified as breeds with "low" teat count (LT), and the other six breeds have a relatively higher teat count (average 15.9), and classified here as breeds with "high" teat count (HT). By applying a similar strategy to identify fixed amino acid substitutions as described previously, we found eleven proteins associated with fixed amino acid substitutions between LT and HT breeds (**Table S9**). We compared the number of amino acids associated with the sites in the two categories of breeds (**Table 3**), and observed that Aspartic acid, Isoleucine, Lysine, Valine, and Tyrosine were associated with the sites more frequently in HT breeds than LT breeds. Our analysis further showed that amino acid substitutions in TAPT1 distinguished the LT/HT pig breeds.

**Table 3.** Number of amino acid substitutions between LT and TT pig breeds.

| Amino Acid | LT | TT | Ratio |
| --- | --- | --- | --- |
| Y | 2 | 9 | 4.5 |
| K | 4 | 17 | 4.3 |
| F | 3 | 11 | 3.7 |
| I | 5 | 13 | 2.6 |
| V | 5 | 12 | 2.4 |
| D | 6 | 12 | 2 |
| N | 5 | 9 | 1.8 |
| E | 7 | 10 | 1.4 |
| Q | 7 | 9 | 1.3 |
| A | 14 | 13 | 0.9 |
| H | 4 | 3 | 0.8 |
| M | 6 | 5 | 0.8 |
| R | 12 | 9 | 0.8 |
| G | 15 | 10 | 0.7 |
| L | 16 | 11 | 0.7 |
| S | 25 | 18 | 0.7 |
| C | 5 | 3 | 0.6 |
| T | 14 | 7 | 0.5 |
| P | 24 | 4 | 0.2 |
| W | 7 | 1 | 0.1 |

### Relevance of placental gene expression to the one-half rule of mammary gland number and litter size

As placental functions are intricately related to the survival of fetuses in the uterus (49–52) and unrelated mammals express common suite of genes in mammary glands and placentae (5), we wanted to explore if placental gene expression is linked to the mammae number and litter size variation between species. We analyzed gene expression data of the term placenta of *Mus musculus, Monodelphis domestica* and *Canis familiaris* (which produce litters and have relatively more teats) and that of *Homo sapiens, Pan paniscus*, and *Loxodonta Africana* (which have fewer teats and produce one offspring at a time). The gene expression data was generated by RNA-seq in a previous study (45). The differential gene expression analysis using the R package *edgeR* (46) identified a small set of genes (n=6) that showed significant changes in placental expression between the two groups (**Figure 9A**). Each of these genes code for proteins associated with the L/NL and T/NT amino acid substitutions identified in the present study. Gene expression of placentae and mammary glands from day-15 pregnant mice was profiled in a recent study in our lab (48). We used these data to compare with genes coding for the proteins associated with the L/NL and T/NT amino acids identified in the present study. This meta-analysis showed that 95.3% of the genes coding the proteins identified in the present study (508 out of 533) were expressed in the placenta as well as in the mammary gland in mice (**Figure 9B)**. This showed that molecular changes associated with the L/NL and T/NT are linked to genes expressed in the placenta suggesting a link of placental function with the molecular basis of the one-half rule.

**Figure 9.** Placental gene expression links mammary gland variation and litter size. (A) Volcano plot showing differential gene expression analysis of placental gene expression data comparing species with high teat number and litter-producing capacity to species with fewer teats and single offspring. The significant genes are indicated. (B) Heatmap showing expression of genes in placenta and mammary gland of day-15 pregnant mice. The scale shows the color codes of gene expression levels.

## Discussion

The results of the present study provide new insights into the evolutionary links between mammary gland location and litter producing capacity in mammals. By analyzing the multiple sequence alignments of protein sequences, we identified 533 proteins that harbored fixed or nearly fixed amino acid substitutions distinguishing litter-producing versus non-litter-producing species (L/NL) as well as the species with thoracic versus non-thoracic mammary gland location (T/NT). Distinct cluster patterns of amino acids in L/NL and T/NT species groups indicated non-random substitution preferences, reflecting constraints related to protein structure or function. Mutual information analyses further demonstrated shared information content among the four species groups, supporting interdependence between amino acid variation associated with mammary gland location and litter size.

The co-occurrence of both substitution types within the same proteins supports the existence of coordinated selective pressures acting on specific protein regions. These genes, based on GO and pathway enrichment analyses, perform a wide variety of different functions; the functions with higher than five-fold enrichment are the epithelial cell apoptotic process, cell adhesion mediated by integrin, regulation of membrane protein ectodomain proteolysis, amine transport, positive regulation of membrane protein ectodomain proteolysis, regulation of gap junction assembly, and sequestering of TGF-beta in extracellular matrix. Several of these functions play crucial roles in the developmental processes of mammary glands, per previous studies (53–57). Moreover, by analyzing the single-cell gene expression data of mouse embryos (44), we found that 300 of the 533 genes identified from the current study are also expressed in the developing mammary gland in mice, suggesting that they are functionally relevant to mammary gland development.

Additionally, it was determined that the majority of these genes were associated with purifying selection (dN/dS < 0.2) suggesting that the observed substitutions are beneficial in nature, possibly for matrotrophy and reproduction of mammals. Evidence for adaptive evolution was observed in a subset of genes (48 of the 533) that exhibited dN/dS ratios greater than 1.0, suggesting that positive selection has acted on specific components of the molecular network linking mammary gland location and reproductive capacity. The presence of adaptive signatures in genes associated with epithelial organization, signaling, and transport is consistent with the dynamic evolutionary pressures acting on mammalian reproductive traits (58, 59). *BayesTraits* analysis further supported correlated evolution between mammary gland location and litter-producing capacity. For the majority of genes analyzed, the dependent model significantly outperformed the independent model, with log Bayes Factors exceeding 2.0, indicating that evolutionary changes in these traits are not independent. This genome-wide signal supports the hypothesis that mammary gland positioning and litter size have evolved in a coordinated manner under shared selective pressures related to reproductive strategy.

The extensive overlap between genes associated with mammary gland variation and those expressed in the placenta suggests a shared molecular basis for prenatal and postnatal maternal investment. Given the essential role of the placenta in fetal development (45, 60–62), coordinated expression of these genes could reflect evolutionary pressures linking placental function with mammary gland traits and litter-producing capacity. Furthermore, our study also identified proteins in which amino acid changes distinguish pig breeds with different teat counts (LT versus HT groups). In pigs, the teat count and their location are important for effectively nursing the piglets. Generally, breeds producing larger litter size have relatively more teats spread throughout the milk line than breeds producing smaller litter size. Moreover, the location of the teats is also important as piglets nursing on the anterior glands gain weight faster than those on the posterior glands (63). We observed that fixed amino acid substitutions distinguishing HT and LT breeds were associated with the protein TAPT1. Importantly, TAPT1 also showed amino acid substitutions in other species analyzed in this study. This convergence suggests that similar molecular mechanisms may underlie both evolutionary divergence and population-level variation in mammary traits. TAPT1 plays a critical role in developmental processes via cilia formation, protein trafficking, and skeletal development, with mutations leading to either lethal or prenatal developmental disorders (64).

Together, our results demonstrate that mammary gland location and litter-producing capacity are evolutionarily coupled through coordinated amino acid substitutions in conserved, developmentally relevant proteins. These findings provide a molecular framework linking mammary gland biology, placental function, and reproductive strategy, and offer new insights into the evolutionary basis of mammalian reproductive diversity.

## Acknowledgements

The authors are thankful to Prof. Thomas B. McFadden for useful discussion about teat number and litter size in pigs.

## Author Contributions

Conception and design: SKB. Data analysis: AB, RM, SKB. Writing and editing: AB, RM, SKB.

## Ethical Statements

No animal experiment was performed in this study

## Data Availability

No experimental data was generated in this study.

## Supplemental Figure Captions

**Figure S1**. A representative multiple sequence alignment showing fixed amino acid substitutions. The gene name and species names are shown. Arrows show position of amino acid substitutions.

**Figure S2**. A bar graph showing the spatial distribution of L/NL and T/NT amino acid substitutions. Distribution of distances between fixed L/NL and T/NT amino acid substitutions within the same proteins, binned in 10–amino acid intervals (shown in x-axis). The frequency of substitutions for each interval is shown the y-axis. The dotted line shows the moving average (n=2) trend line.

**Figure S3**. Comparison between sample dendrogram patterns. Left: comparison of cophenetic distance (y-axis) between clusters in pairwise manner between species groups (x-axis). Right: Cluster correlation patterns between species groups. The scale on the right shows color codes for correlation levels.

