## Supplemental Text 1 for "Co-occurring Amino Acid Substitutions Reveal Shared Evolutionary Links between Mammary Gland Location and Litter Size in Mammals"

**Text S1. Genome assembly versions of the species used in the analysis.**

| Animal | Species name | Genome assembly used in the analysis |
| --- | --- | --- |
| Whale | *Delphinapterus leucas* | Delphinapterus_leucas.ASM228892v3 |
| Horse | *Equus caballus* | Equus_caballus.EquCab3.0 |
| Gorilla | *Gorilla gorilla* | Gorilla_gorilla.gorGor4 |
| Human | *Homo sapiens* | Homo_sapiens.GRCh38 |
| Elephant | *Loxodonta africana* | Loxodonta_africana.loxAfr3 |
| Gibbon | *Nomascus leucogenys* | Nomascus_leucogenys.Nleu_3.0 |
| Chimpanzee | *Pan troglodytes* | Pan_troglodytes.Pan_tro_3.0 |
| Orangutan | *Pongo abelii* | Pongo_abelii.Susie_PABv2 |
| Cow | *Bos taurus* | Bos_taurus.ARS-UCD2.0 |
| Cat | *Felis catus* | Felis_catus.F.catus_Fca126_mat1.0 |
| Ferret | *Mustela putorius furo* | Mustela_putorius_furo.MusPutFur1.0 |
| Squirrel | *Sciurus vulgaris* | Sciurus_vulgaris.mSciVul1.1 |
| Dog | *Canis lupus familiaris* | Canis_lupus_familiaris.ROS_Cfam_1.0 |
| Mouse | *Mus musculus* | Mus_musculus.GRCm39 |
| Rabbit | *Oryctolagus cuniculus* | Oryctolagus_cuniculus.OryCun2.0 |
| Fox | *Vulpes vulpes* | Vulpes_vulpes.VulVul2.2 |
| Rat | *Rattus norvegicus* | Rattus_norvegicus.GRCr8 |
| Pig | *Sus scrofa* | Sus_scrofa.Sscrofa11.1 |
