## Supplemental Text 2 for "Co-occurring Amino Acid Substitutions Reveal Shared Evolutionary Links between Mammary Gland Location and Litter Size in Mammals"

Text S2. Extracting protein sequences commonly present across the selected species, and performing multiple sequence alignment for each protein.


# Trim the fast header to keep gene symbol only (assuming that the current folder contains the fasta files of protein sequences from each species that were obtained from Ensembl)

for i in $(ls *.fa); do sed 's/^.*gene_symbol:/>/' $i | awk '{print $1}' > $i'_gene_symbol_fasta'; done


# Convert multiline fasta to single-line fasta

for i in $(ls *_gene_symbol_fasta); do awk '{if(NR==1) {print $0} else {if($0 ~ /^>/) {print "\n"$0} else {printf $0}}}' $i > $i'_sequences_singlline_fasta'; done


# Convert single-line fasta to tabular form

for i in $(ls *_sequences_singlline_fasta); do awk -v OFS="\t" 'BEGIN{RS=">"}{gsub("\n","\t",$0); gsub(">","",$0); print $0}' $i > $i'_gene_sequence_table';done


# Extract the longest protein sequence for each gene
for i in $(ls *gene_sequence_table); do awk '{print $1, $2, sprintf("%s", length($2))}' $i | sort -k1,1 -k3,3nr | sort -u -k1,1  | awk -v OFS='\t'  '{print $1, $2}' > $i'_longest_sequence'; done

# Concatenate the files into a single file

awk '{print $0, FILENAME}' *.pep.all.fa_gene_symbol_fasta_sequences_singlline_fasta_gene_sequence_table_longest_sequence* | sed 's/.pep.all.fa_gene_symbol_fasta_sequences_singlline_fasta_gene_sequence_table_longest_sequence//g' > all_longst_protein_id_seq_species


# Retain the sequences of the proteins commonly present across the 18 species, and remove those are not.

awk 'NR==FNR{a[$1]++; next} a[$1]==18' all_longst_protein_id_seq_species all_longst_protein_id_seq_species > common_gene_sequences

# Creating the individual fasta files for of the common proteins

for i in $(awk  '{print $7}' common_gene_sequences | uniq);  do grep -w $i common_gene_sequences |  awk  -v OFS='\t' '{print $1"_"$4, $2}'  |  awk  -v OFS='\n' '{$1 = ">" $1} 1' > "species18_seq_"$i; done
