## Supplemental Text 3 for "Co-occurring Amino Acid Substitutions Reveal Shared Evolutionary Links between Mammary Gland Location and Litter Size in Mammals"

Text S3. Performing multiple sequence alignment of amino acid sequences of each protein among the 18 species, and identification of sites that are fixed between L vs NL or T vs NT groups.

# Generate multiple  alignments of sequences for each gene using cluster omega (assumes clustalo is installed and accessible).

for i in $(ls species18_seq_*); do clustalo -i $i -o aln_$i -v; done

# Convert aligned multiline fasta sequences to single line fasta
for i in $(ls aln_species18_seq_*); do awk '{if(NR==1) {print $0} else {if($0 ~ /^>/) {print "\n"$0} else {printf $0}}}' $i > $i'_sequences_singlline_fasta'; done

# Convert single-line fasta to tabular form
for i in $(ls *_sequences_singlline_fasta); do awk '/^>/ {printf "%s\t", substr($0, 2)} !/^>/ {print $0}' $i > $i'_gene_sequence_table';done

# Concatenate the files to one file
awk '{print $0, FILENAME}' *_gene_sequence_table | sed 's/aln_species18_seq_//g' | sed 's/_sequences_singlline_fasta_gene_sequence_table//g' | sed 's/_/ /g' > ALL_ALN_TABLE_FORM

# Extract aligned sequences from the combined file individual groups (L, NL, T and NT)

grep -E 'Dog|Cat|Mouse|Ferret|Rabbit|Rat|Squirrel|Pig|Fox' ALL_ALN_TABLE_FORM > litter

grep -E 'Cow|Whale|Horse|Gorilla|Human|Elephant|Gibbon|Chimpanzee|Orangutan' ALL_ALN_TABLE_FORM > nonlitter

grep -E 'Gorilla|Human|Elephant|Gibbon|Chimpanzee|Orangutan' ALL_ALN_TABLE_FORM > thoracic

grep -E 'Cow|Dog|Whale|Horse|Cat|Mouse|Ferret|Rabbit|Rat|Squirrel|Pig|Fox' ALL_ALN_TABLE_FORM > nonthracic

# Identify variant sites and count their frequency (how many species in the group are associated with the variant amino acids). This is done for each of 4 groups.


# L group
awk '{gene=$4; alignment=$3; for(i=1; i<=length(alignment); i++) {aa=substr(alignment, i, 1); counts[gene,i,aa]++}} END {for(key in counts) {split(key, arr, SUBSEP); print arr[1], arr[2], arr[3], counts[key]}}' litter > counts_L

# NL group
awk '{gene=$4; alignment=$3; for(i=1; i<=length(alignment); i++) {aa=substr(alignment, i, 1); counts[gene,i,aa]++}} END {for(key in counts) {split(key, arr, SUBSEP); print arr[1], arr[2], arr[3], counts[key]}}' nonlitter > counts_NL


# T group

awk '{gene=$4; alignment=$3; for(i=1; i<=length(alignment); i++) {aa=substr(alignment, i, 1); counts[gene,i,aa]++}} END {for(key in counts) {split(key, arr, SUBSEP); print arr[1], arr[2], arr[3], counts[key]}}' ALL_ALN_THORASIC > counts_T


# NT group

awk '{gene=$4; alignment=$3; for(i=1; i<=length(alignment); i++) {aa=substr(alignment, i, 1); counts[gene,i,aa]++}} END {for(key in counts) {split(key, arr, SUBSEP); print arr[1], arr[2], arr[3], counts[key]}}' ALL_ALN_NONTHORASIC > counts_NT


# Identify variant sites that are fixed or nearly fixed between species groups (see main text for explanation).

# Identify sites that are fixed in all species within the groups

awk '$4 == 9' counts_L |sort > counts9_L

awk '$4 == 9' counts_NL |sort > counts9_NL

awk '$4 == 6' counts_T |sort > counts6_T

awk '$4 == 6' counts_NT |sort > counts6_NT


awk '$4 == 8' counts_L |sort > counts8_L

awk '$4 == 8' counts_NL |sort > counts8_NL

awk '$4 == 5' counts_T |sort > counts5_T

awk '$4 == 5' counts_NT |sort > counts5_NT


awk 'NR==FNR{a[$1,$2]=$0; next} ($1,$2) in a{print a[$1,$2], $0}' counts9_L counts9_NL | awk '$3 != $7 {print}' > fixed_all_L/NL

awk 'NR==FNR{a[$1,$2]=$0; next} ($1,$2) in a{print a[$1,$2], $0}' counts9_L counts8_NL | awk '$3 != $7 {print}' > fixed_all_L_most_NL


awk 'NR==FNR{a[$1,$2]=$0; next} ($1,$2) in a{print a[$1,$2], $0}' counts8_L counts9_NL | awk '$3 != $7 {print}' > fixed_most_L_all_NL

# Identify sites that are either fixed in all or most species between T and NT species groups

awk 'NR==FNR{a[$1,$2]=$0; next} ($1,$2) in a{print a[$1,$2], $0}' counts6_T counts12_NT | awk '$3 != $7 {print}' > fixed_all_T/NT

awk 'NR==FNR{a[$1,$2]=$0; next} ($1,$2) in a{print a[$1,$2], $0}' counts6_T counts11_NT | awk '$3 != $7 {print}' > fixed_all_T_most_NT


awk 'NR==FNR{a[$1,$2]=$0; next} ($1,$2) in a{print a[$1,$2], $0}' counts5_ counts12_NT | awk '$3 != $7 {print}' > fixed_most_T_all_NT
