## Supplemental Figure 1 for "Co-occurring Amino Acid Substitutions Reveal Shared Evolutionary Links between Mammary Gland Location and Litter Size in Mammals"

```
                      1        10        20        30        40
                      |--------+---------+---------+---------|
PLA2G4A_Cow           YWGSAFSILFNRVLGYSGSQSKGSTMEEELENITAKHIYS
PLA2G4A_Whale         YWGSAFSILFNRVLGYSGSQSKGSTMEEELENITAKHIYS
PLA2G4A_Dog           YWGSAFSILFNRVLGYSGSQNKGSTMEEELENITAEHIYS
PLA2G4A_Cat           YWGSAFSILFNRVLGYSGSQNKGSTMEEELENITAEHIYS
PLA2G4A_Fox           YWGSAFSILFNRVLGYSGSQNKGSTMEEELENITAEHIYS
PLA2G4A_Ferret        YWGSAFSILFNRVLGYSGSQNKGSTMEEELENITAEHIYS
PLA2G4A_Horse         YWGSAFSILFNRVLGYSGSQNKGSTMEEELENITAKHIYS
PLA2G4A_Pig           YWGSAFSILFNRVLGYSGSQNKGSTMEEELENITAKHIYS
PLA2G4A_Squirrel      YWGSAFSILFNRVLGYSGSQNKGSTMEEELENITAKHIYS
Pla2g4a_Rat           YWGSAFSILFNRVLGYSGSQNKGSTMEEELENITAKHIYS
Pla2g4a_Mouse         YWGSAFSILFNRVLGYSGSQNKGSTMEEELENITAKHIYS
PLA2G4A_Rabbit        YWGSAFSILFNRVLGYSGSHNKGSTMEEELENITAKHIYS
PLA2G4A_Elephant      YWGSAFSILFNRVLGYSASQSKGSTMEEELENITAKHIYS
PLA2G4A_Gorilla       YWGSAFSILFNRVLGYSGSQSRGSTMEEELENITTKHIYS
PLA2G4A_Orangutan     YWGSAFSILFNRVLGYSGSQSRGSTMEEELENITTKHIYS
PLA2G4A_Gibbon        YWGSAFSILFNRVLGYSGSQSRGSTMEEELENITTKHIYS
PLA2G4A_Human         YWGSAFSILFNRVLGYSGSQSRGSTMEEELENITTKHIYS
PLA2G4A_Chimpanzee    YWGSAFSILFNRVLGYSGSQSRGSTMEEELENITTKHIYS
```
