## Supplementary figures and images for "Co-occurring Amino Acid Substitutions Reveal Shared Evolutionary Links between Mammary Gland Location and Litter Size in Mammals"

### Supplemental Figure 2

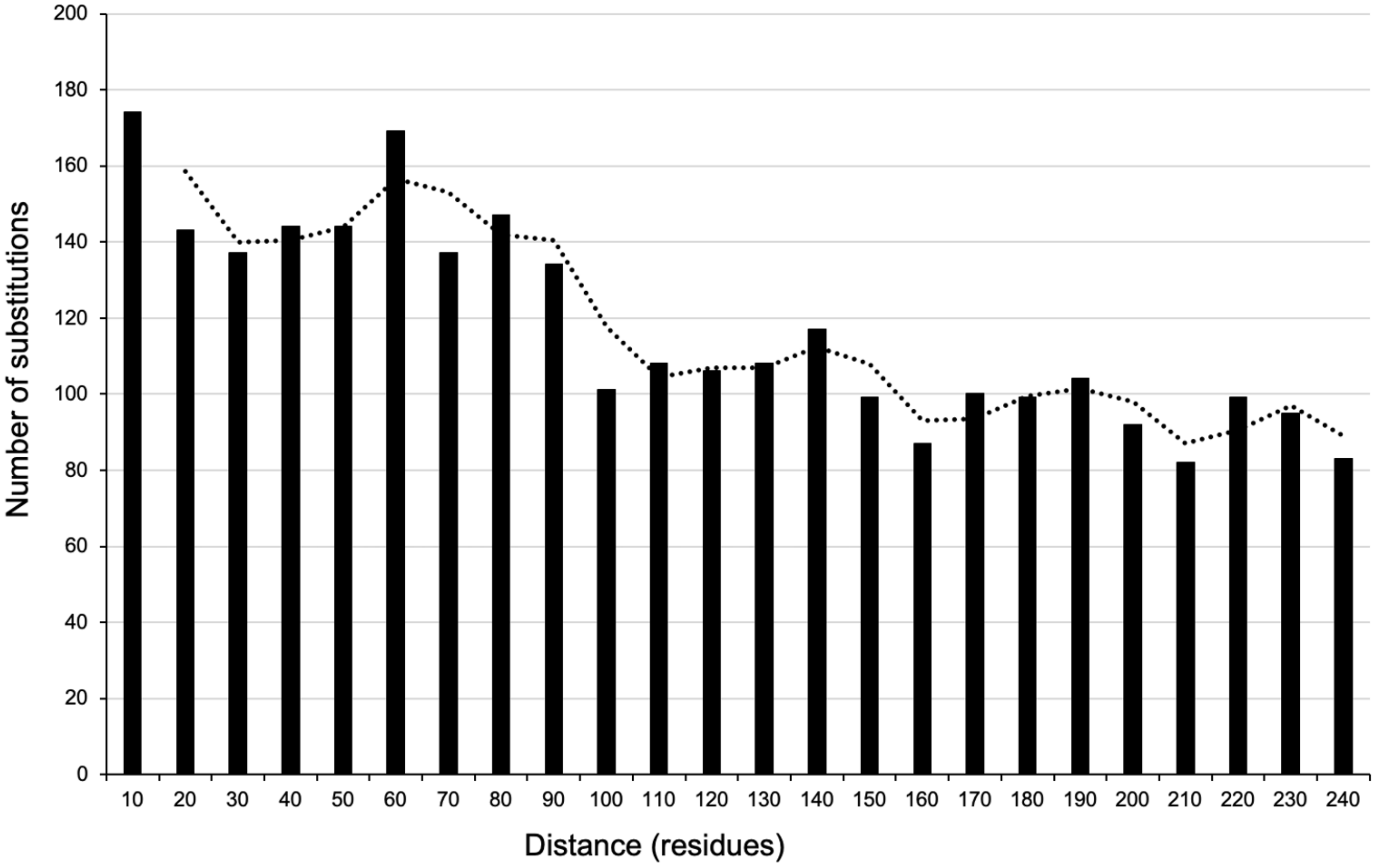

### Supplemental Figure 3

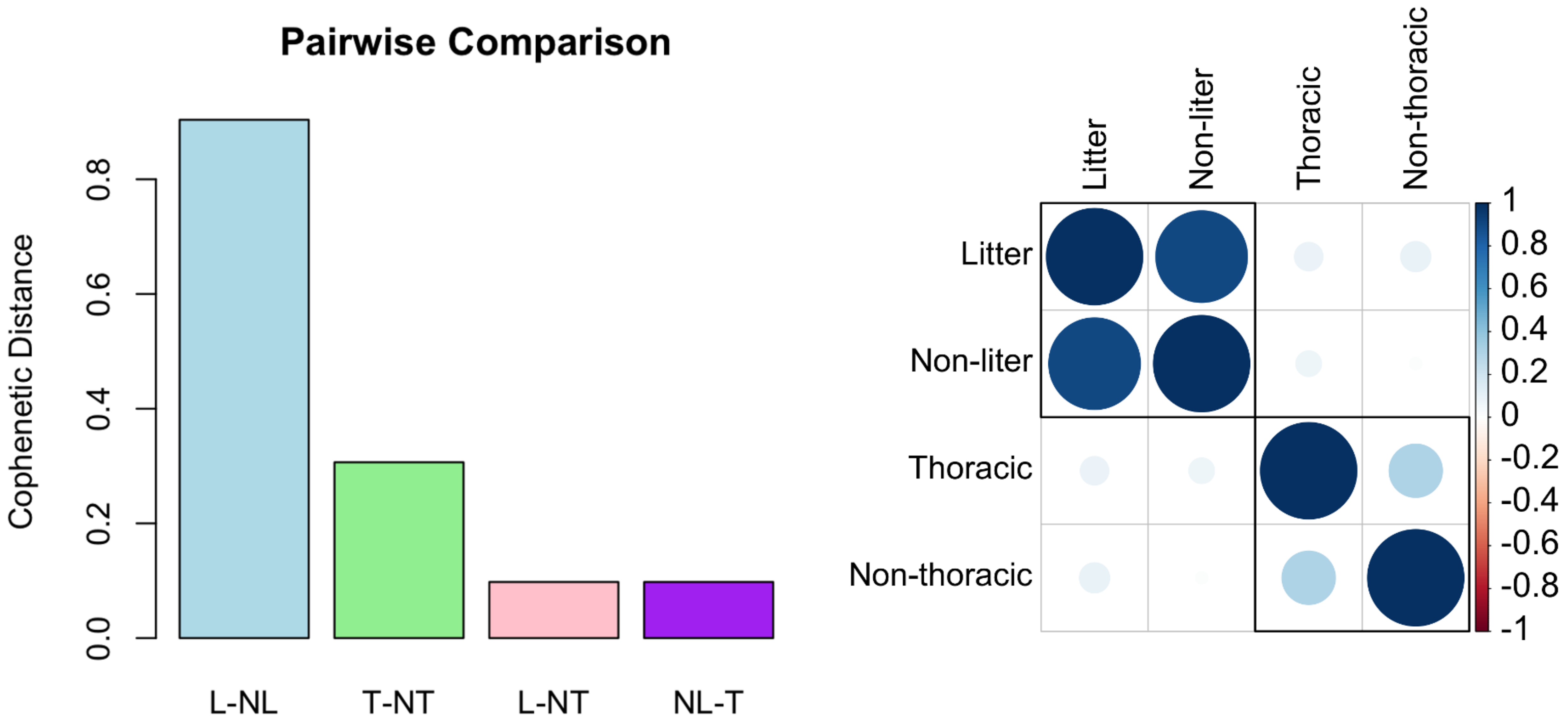
